# *Drosophila* males integrate song and pheromones using context-specific strategies

**DOI:** 10.1101/2025.07.25.666755

**Authors:** Adrián Palacios-Muñoz, Jan Clemens

## Abstract

Animals need to integrate sensory information from multiple modalities to interact with their environment and with others. In male *Drosophila*, multiple sensory cues modulate social behaviors such as courtship and aggression, but the strategies by which multimodal cues are combined to inform social behavior are not known. Here, we focus on how courtship song and taste cues are integrated to shape the social behavior of male flies. By combining cue manipulations with computational modeling, we assessed the individual and combined contributions of song and taste. Our results reveal three context-specific integration strategies. Overall social interactions were driven by a linear combination of song and taste. Aggression was driven by song, independent of taste cues. Courtship was controlled by a nonlinear integration of song and female taste cues, with female taste flipping the valence of song from suppressing to enhancing courtship. Our results show how context-specific integration strategies allow animals to recognize social scenarios and produce flexible, context-appropriate behaviors.

## Introduction

Behavioral decisions often rely on multimodal cues. We assess a fruit’s ripeness by combining visual, tactile, and olfactory cues—the perfect tomato should be deep red, not too soft or too firm, and smell aromatically. Just as multimodal integration helps identify a perfect tomato, it also plays a key role in social interactions, helping animals recognize individuals, assess competition, and find a suitable mate. For instance, we evaluate facial expressions alongside speech and can detect dishonesty when cues conflict (Zuckerman and Neeb, 1979; McGurk and MacDonald, 1976). Similarly, birds, frogs, and insects integrate multiple sensory cues to guide their social interactions (Nowlan et al., 2025; McRae et al., 2023; Rolland et al., 2022; Fleischer et al., 2022; Auer and Benton, 2016; Krstic et al., 2009; Dickson, 2008).

To be efficient, multimodal integration should be context-specific, ignoring irrelevant or distracting cues, and changing the valence of one cue based on the presence of another cue (Alais and Burr, 2004; Körding et al., 2007; Cao et al., 2019). Studying multimodal integration during naturalistic social interactions is challenging, because of the complexity of the behavioral readouts and the difficulty of controlling social cues. To generate testable hypotheses about the underlying neural mechanisms, it is essential to combine quantitative behavioral analysis with targeted manipulation of multimodal stimuli (Colonius and Diederich, 2017).

Here, we study multimodal integration during social behavior of male *Drosophila melanogaster*, leveraging their robust, well-characterized behaviors and the ease of manipulating their sensory perception. Male fruit flies rely on multiple modalities—acoustic, mechanosensory, visual, olfactory, taste—to identify social targets and select target-appropriate behaviors, such as courtship or aggression. Here, we focus on the integration of taste and acoustic cues, as they individually have robust and sometimes opposing effects on social behavior.

Taste cues—cuticular hydrocarbons—are sampled by foreleg tapping and provide information about the sex, species, and mating status of another fly (Rybak et al., 2002; Fan et al., 2013; Seeholzer et al., 2018; Yew and Chung, 2015). Female-specific taste cues (e.g., 7,11-dienes) are detected by neurons in the foreleg tarsi co-expressing ppk23 and ppk25 channels (Clowney et al., 2015; Kallman et al., 2015), and promote male courtship (Antony and Jallon, 1982; Grillet et al., 2006; Foley et al., 2007; Billeter et al., 2009; Kohl et al., 2015). Male-specific taste cues (e.g., 7-T) are detected by neurons expressing ppk23 but not ppk25 channels, as well as other gustatory receptor neurons (GRNs) such as Gr32a (Fan et al., 2013; Clowney et al., 2015; Kallman et al., 2015), and can suppress courtship (Seeholzer et al., 2018; Fan et al., 2013; Wang et al., 2011; Yew et al., 2009) or trigger aggression (Fernández et al., 2010).

Acoustic cues are produced by extending and vibrating one or both wings. Courtship song, typically directed at females, is produced by unilateral wing vibration and consists of two main modes: pulse song, trains of pulses with an interval of 30–40 ms, and sine song, a continuous oscillation around 150 Hz (Ewing and Bennet-Clark, 1968; Rybak et al., 2002). Agonistic song, typically directed at males, is produced by bilateral wing vibration and consists of longer, irregular pulses (Versteven et al., 2017; Jonsson et al., 2011; Hindmarsh Sten et al., 2025).

Here, we focus on courtship song because it can evoke courtship or aggression, depending on the behavioral context. Notably, hearing song can drive male-male courtship in groups of males, suggesting that it may override the inhibitory effect of the otherwise suppressive male taste cues (Crossley et al., 1995; Yoon et al., 2013). However, this interaction has not been tested explicitly. When a female is present, song drives aggression rather than courtship between males (Versteven et al., 2017; Hindmarsh Sten et al., 2025), suggesting that its effect in males may depend on additional cues such as taste.

While central neurons that process either song or taste cues to drive courtship or aggression have been identified in the fly brain (Baker et al., 2022; Clowney et al., 2015; Kallman et al., 2015; Hindmarsh Sten et al., 2025; Zhou et al., 2014; Zhou et al., 2015; Asahina et al., 2014; Shankar et al., 2015), how song-taste integration is implemented in the brain is not known.

Here, to examine how male *Drosophila* integrate taste cues and courtship song to guide social behavior, we combine manipulations of taste perception and courtship song with quantitative behavioral modeling. We find that males integrate song and taste cues in three distinct context-specific strategies: (1) linear integration to modulate male- and female-directed interactions, (2) song-driven male-male aggression independent of taste cues, and (3) a switch in the behavioral valence of song by female taste to promote of courtship.

By revealing diverse strategies of sensory cue integration, we provide new insights into flexible, context-specific multisensory integration, and pave the way for identifying the neural circuits underlying these computations.

## Results

### Song and male taste cues are linearly integrated to modulate male-male interactions

To determine how male fruit flies integrate taste and courtship song during social interactions, we first manipulated taste and acoustic cues and measured their effects on male-male interactions. We recorded pairs of wild-type males for 30 minutes (Fig. 1A) and quantified their interactions as the fraction of time spent within 4 mm of each other (*social interaction index*). To distinguish social interactions from time spent passively in proximity, we only counted as interactions periods during which at least one fly moved faster than 1 mm/s.

**Figure 1:**
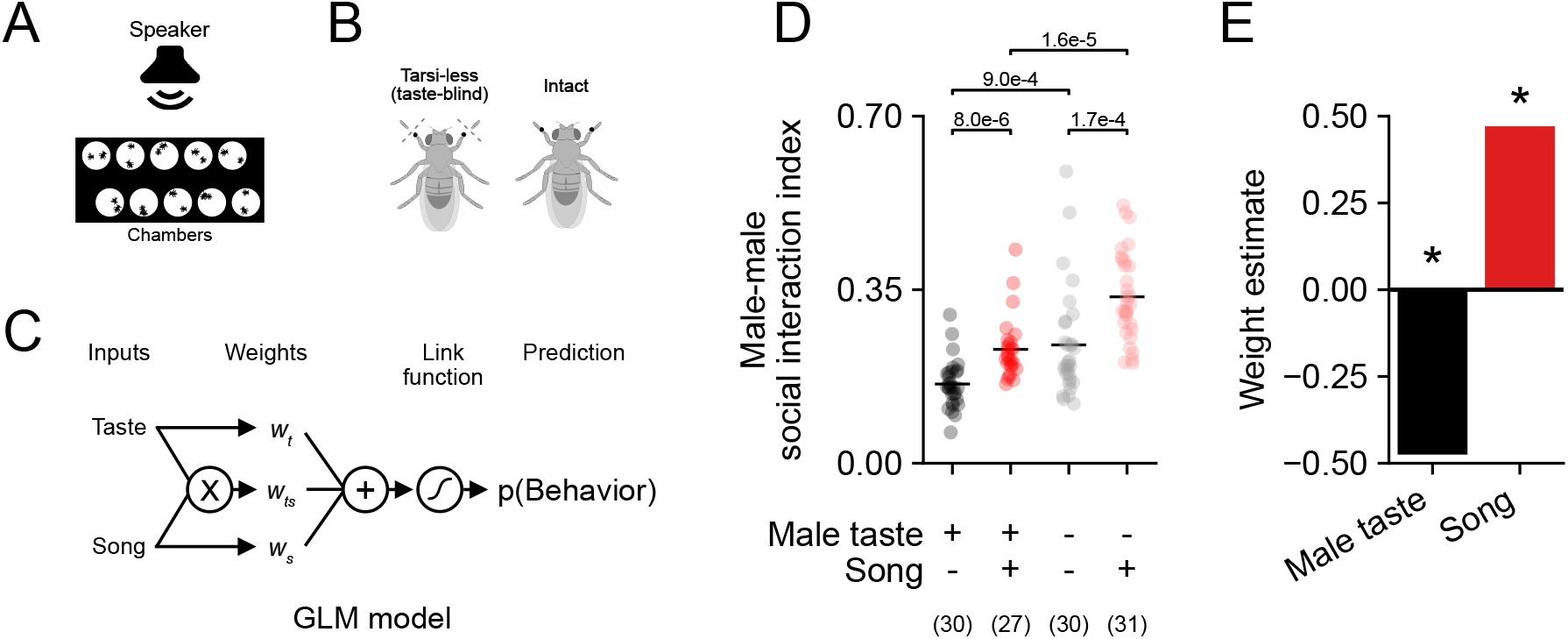
Song and male taste cues are linearly integrated to control male-directed social interactions. **A** Experimental setup. Flies were recorded in a multi-arena chamber, illuminated from below with infrared light, and recorded using a top-down camera. A loudspeaker positioned behind the holder delivered song stimuli. **B** Sensory manipulations. Foreleg tarsi were removed one day before the experiment under cold anesthesia. Intact (sham-treated) males were anesthetized and handled identically but without tarsi removal. **C** Structure of the Generalized Linear Models (GLMs). Inputs are weighted individually (song: *w*^*s*^ taste: *w*^*t*^) and linearly summed before passing the result through a logistic function to predict the social interaction index. Nonlinear integration was accounted for by a cue interaction term, *w*^*ts*^. **D** Social interaction index for male-male pairs. Song playback (red) and taste manipulation increased social interactions. Dots correspond to individual male-male pairs (N provided below the panel). Black dots correspond to experiments with no playback. For pair-wise comparisons, we tested each group for normality with a Shapiro test, and conducted the comparison with an independent t-tests for unequal variances if both groups were normally distributed, or with a two-sided Mann-Whitney U test otherwise. All p-values are reported after Bonferroni correction for multiple comparisons. **E** Weights from the best GLM (linear vs null: *p* = *3 .7 e*^*−14*^, one-sided test of the hypothesis that linear reduces deviance more than the null model, nonlinear vs linear: *p* = *0 .69*) predicting male-male interactions in D. Male taste cues have a negative weight (*w*^*t*^ = *− 0 .48, p* = *3 .5 e*^*−10*^, black) and song has a positive weight (*w*^*s*^ = *0 .47, p* = *3 .8 e*^*−10*^, red). This indicates that perceiving male taste suppresses and playback of song promotes social interaction between males. Asterisk above bar indicate significance of weight in the model. aggression independently of taste cues. By contrast, song’s effect on male-directed courtship was much smaller and individually non-significant (Fig. 2B).

We eliminated the males’ ability to detect suppressive male taste cues via foreleg tapping by removing their foreleg tarsi. This manipulation increased interactions by 49% relative to baseline (Fig. 1D). Similarly, playing back conspecific pulse song (“song”) for 20 s every 2 minutes promoted interactions by 44%. These results confirm the previously reported effects: Male taste cues suppress, and song promotes, male-male interactions (Seeholzer et al., 2018; Fan et al., 2013; Wang et al., 2011; Yew et al., 2009; Crossley et al., 1995; Versteven et al., 2017; Yoon et al., 2013). To test the integration of taste and song, we combined song playback with the removal of foreleg tarsi. This increased interactions by 110%, approximately the sum of the individual effects, suggesting that male taste cues and song are linearly integrated.

To quantitatively test between linear and nonlinear modes of song-taste integration, we fitted Generalized Linear Models (GLMs) to the data (Fig. 1C). The GLMs assign weights to taste and song cues, sum them, and apply a logistic function to predict the social interaction index. The magnitude and sign of each cue’s weight indicate its importance and valence during social interactions. A linear integration model includes only weights for the individual cues, taste and song. A model of nonlinear integration contains an additional interaction term, which allows the effect of one cue to be changed by the presence of the other. We compared the ability of linear and non-linear integration models to account for the behavior using the Akaike Information Criterion (AIC, Akaike, 1974), which balances model performance with model complexity (the number of weight parameters).

The linear model best predicts the data in the above experiments, with no evidence for nonlinear integration (Fig. 1E, see Table S1-S2 for model selection, parameters and statistics). Song and taste had equal but opposite weights (*w*_*s*_ = *0 .47, w*_*t*_ = *− 0 .48*), consistent with our observation that song fully cancels the suppressive effect of male taste cues.

### Song, but not taste, promotes male-directed aggression

Thus far, we have shown that song and taste cues are independently and linearly integrated to modulate social interactions between two males. However, previous studies have reported different effects of the individual cues depending on the type of interaction: Male taste suppresses courtship and promotes aggression, while song promotes both courtship and aggression (Schilcher, 1976; Ejima and Griffith, 2008; Rybak et al., 2002; Deutsch et al., 2019; Yoon et al., 2013; Hindmarsh Sten et al., 2025; Li et al., 2018). To test how both cues are integrated to control these behaviors, we analyzed the data from the previous experiments, now distinguishing between courtship and aggression.

We identified courtship and aggression based on wing movement, respectively: During courtship, males produce courtship song using unilateral wing extensions (UWEs) (Clemens et al., 2018; Yamamoto and Koganezawa, 2013; BENNET-CLARK and EWING, 1967). Aggression, is associated with agonistic song produced by brief bilateral wing extensions (BWEs) (Jonsson et al., 2011; Hindmarsh Sten et al., 2025; Versteven et al., 2017; Kravitz and Fernandez, 2015; Duistermars et al., 2018).

We first confirmed that song drives aggression in male pairs in our assay, indicated by an increase in BWEs (Hindmarsh Sten et al., 2025). Since aggression functions to interrupt rival courtship behavior, male taste cues may be crucial for detecting competitors and triggering aggressive responses. Alternatively, male taste cues may be dispensable for this form of aggression, as males may need to repel rivals before touching them. Indeed, exposure to courtship song increased male aggression, independent of the ability of males to taste their interaction partner (Fig. 2A left). Accordingly, the amount of aggression between two males was best explained by a linear integration model, with a strong positive weight for song (*w*_*s*_ = *1 .78, p* = *3 .5 e*^*−16*^) and weak and non-significant weight for taste (Fig. 2A right). These results show that song drives

**Figure 2:**
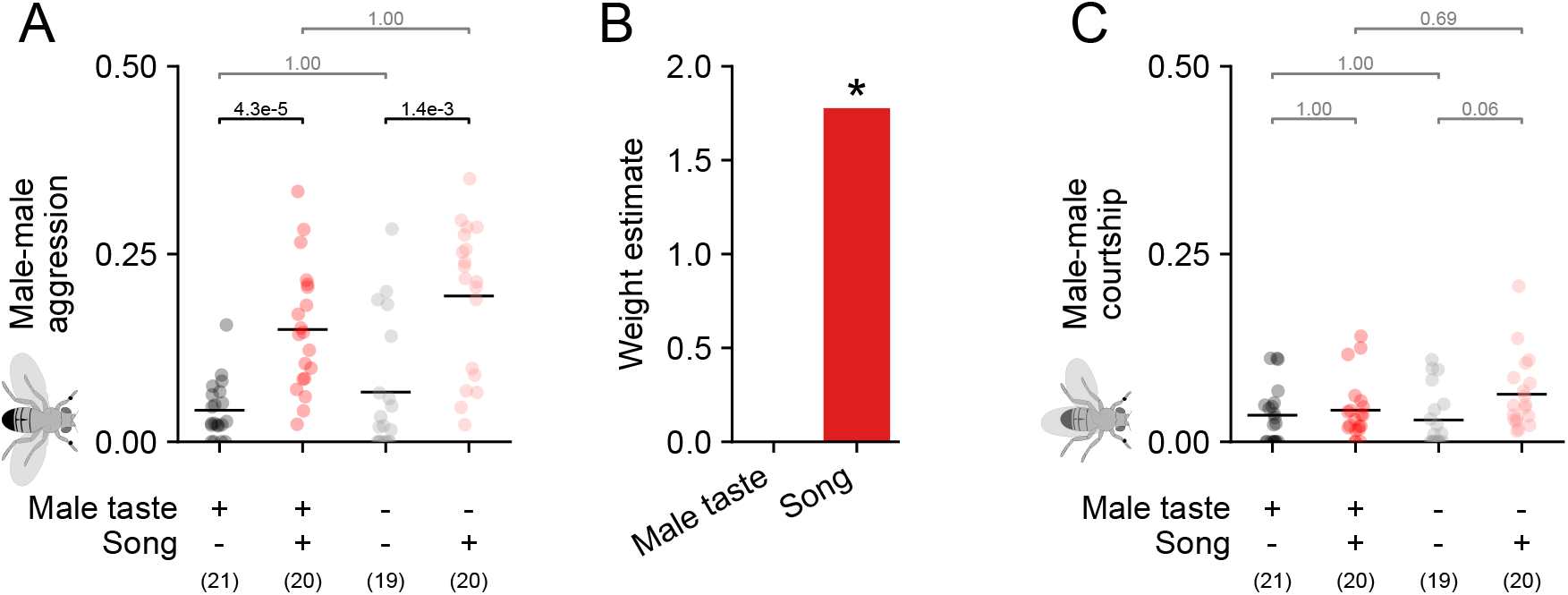
Song increases male-directed aggression independent of taste cues. **A** Probability of aggression (bilateral wing extension) during social interactions for taste and song (red) manipulations in pairs of males. **B** Weights from the selected GLM (linear vs. null: *p* = *1 .5 e*^*−12*^, nonlinear vs. linear: *p* = *0 .16*) predicting male-male aggression data from A (see tables S1 and S2 for details). Taste has a small and nonsignificant weight (*w*^*t*^ = *− 0 .02, p* = *0 .92*, black), indicating that interfering with the detection of male taste cues does not affect aggression. Song has a strong and positive weight (*w*^*s*^ = *1 .78, p* = *3 .5 e*^*−16*^, red), indicating that song promotes aggression. Asterisk above bar indicate significance of weight in the model. **C** Probability of courtship (unilateral wing extension) during social interactions for taste and song (red) manipulations in pairs of males. Model fit omitted due to lack of significant effects. Dots in A and C correspond to individual male-male pairs (N provided below the panel). For pair-wise comparisons, we tested each group for normality with a Shapiro test, and conducted the comparison with an independent t-tests (song groups in A) for unequal variances if both groups were normally distributed, or with a two-sided Mann-Whitney U test otherwise (all other tests). All p-values are reported after Bonferroni correction for multiple comparisons.

In summary, courtship song and male taste cues are linearly integrated with opposing signs to modulate the males decision to interact with another male. Neither song nor taste strongly affected courtship in pairs of males, suggesting that previously reported courtship promoting effects of song rely on the group context (Yoon et al., 2013). By contrast, song alone was sufficient to drive aggression in male pairs, with no contribution from taste. Overall, these results show that song and taste cues are integrated in diverse ways to control general social interaction, courtship, and aggression in male-male interactions.

### Song and female taste cues are linearly integrated to control social interaction

Unlike male taste cues, female taste cues are known to promote courtship (Bray and Amrein, 2003; Yamamoto and Koganezawa, 2013; Clowney et al., 2015). To examine how males integrate song with female taste cues, we manipulated the perception of taste cues and played back courtship song as before, but now in males paired with mature virgin females. To isolate the effect of song on male behavior, we deafened the female partners by removing their aristae one day before the experiments.

We find that males increased female-directed interactions when able to acquire taste cues through tapping (Fig. 3A). Playback of song increased interactions only when males were prevented from detecting female taste cues. When males could detect female taste cues, their interaction rates were already high, leaving little room for further increases with song. Removal of taste cues reduced the baseline and revealed the song effect.

**Figure 3:**
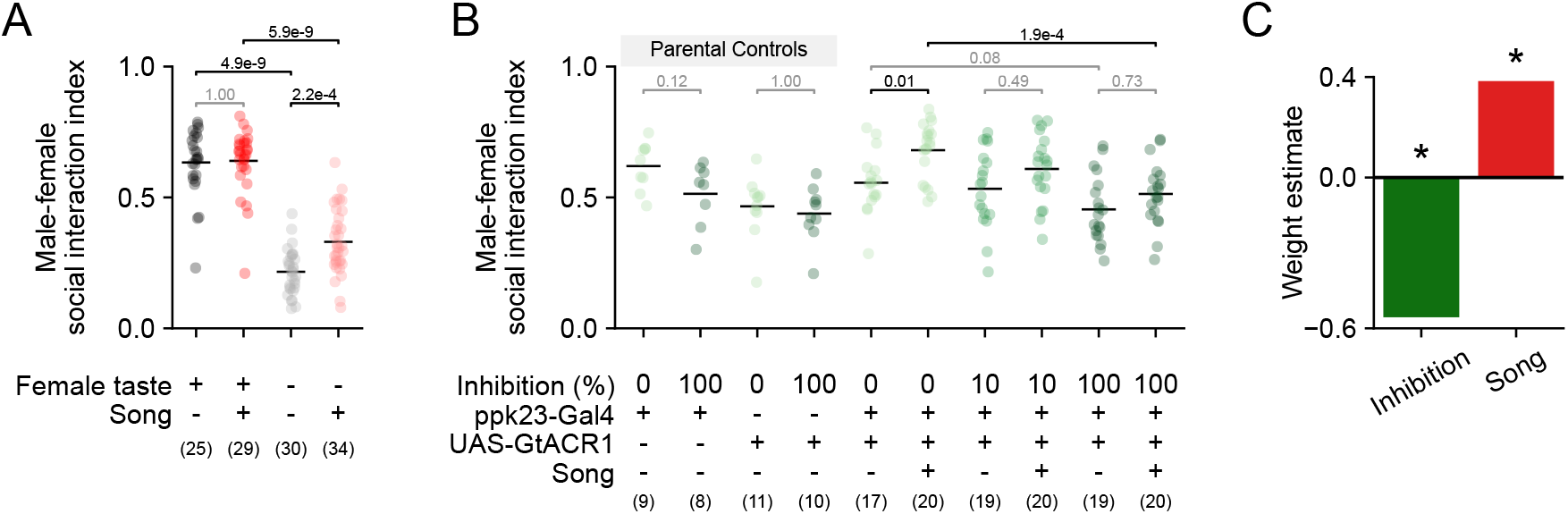
Song and female taste cues are linearly integrated to control female-directed social interactions. **A** Social interaction index for wild-type male-female pairs. Removing taste cues strongly decreased social interactions. Playback of song (red) increased social interactions in males where taste was manipulated. **B** Effect of ppk23 inhibition on male-female social interactions. Social interaction index for parental controls and ppk23-inhibited males. No significant effect was found in controls, confirming that LED exposure alone did not introduce confounding effects. Song significantly increased social interaction in group without inhibition (0% inhibition, no green LED exposure). Inhibition of taste receptors significantly decreased social interaction. **C** Weights from the selected GLM (linear vs. null: *p* = *6 .9 e*^*−9*^, nonlinear vs. linear: *p* = *0 .74*) predicting the social interaction index data from B (see tables S3 and S4 for details). Inhibition is associated with a negative weight (*w*_*inh*_ = *− 0 .56, p* = *2 .2 e*^*−7*^, green) indicating that interfering with the perception of female taste cues reduces social interactions. Song is associated with a positive weight (*w*^*s*^ = *0 .38, p* = *5 .7 e*^*−5*^, red) and this promotes social interactions. Asterisk above bar indicate significance of weight in the model. Dots in A and B correspond to individual male-male pairs (N provided below the panel).Green darkness in B encode the inhibition strength, as indicated below each group. For pair-wise comparisons, we tested each group for normality with a Shapiro test, and conducted the comparison with an independent t-tests for unequal variances if both groups were normally distributed, or with a two-sided Mann-Whitney U test otherwise (intact wild-type groups, first two groups from panel A). All p-values are reported after Bonferroni correction for multiple comparisons.

To avoid this saturation effect, we gradually reduced taste inputs using optogenetic inhibition of foreleg taste receptors, rather than fully removing the tarsal segment. We expressed the inhibitory channel rhodopsin GtACR1 (Mauss et al., 2017) in the ppk23-expressing neurons (ppk23-GtACR1 males), which label taste receptor neurons specifically sensitive to male and female pheromones (Toda et al., 2012; Kallman et al., 2015; Clowney et al., 2015).

In ppk23-GtACR1 males, increasing light intensity reduced interactions with females compared to parental controls (Fig. 3B). With intact taste receptors (0% inhibition), song significantly increased interactions. This effect decreased with increasing inhibition of taste receptors. Accordingly, the data were best explained by a linear integration model, with a positive weight for song (*w*_*s*_ = *0 .38, p* = *5 .7 e*^*− 5*^) and a negative weight for taste inhibition (*w*_*inh*_ = *− 0 .56, p* = *2 .2 e*^*− 7*^) (Fig. 3C).

### Song-driven increase in interaction is feature-specific

To test whether the effect of song on female-directed interactions was specific to conspecific pulse song, we tested song-taste integration using manipulated song stimuli. In addition to conspecific pulse song with a 36 ms inter-pulse interval (IPI), we also tested modified pulse song with shorter (16 ms) and longer (96 ms) IPIs. Song detector neurons in the male brain respond less to these modified stimuli and should therefore be less effective in driving interactions (Deutsch et al., 2019; Yoon et al., 2013; Li et al., 2018; Zhou et al., 2015).

While exposing male-female pairs to songs with different IPIs, we inhibited the ppk23 neurons at different levels. When taste perception was intact (0% inhibition), social interactions showed the expected band-pass tuning for IPI, with a peak at 36 ms (Fig. 4A). This peak flattened with increasing inhibition, resulting in a loss of IPI tuning, consistent with the absence of song effects observed above (Fig. 3B).

**Figure 4:**
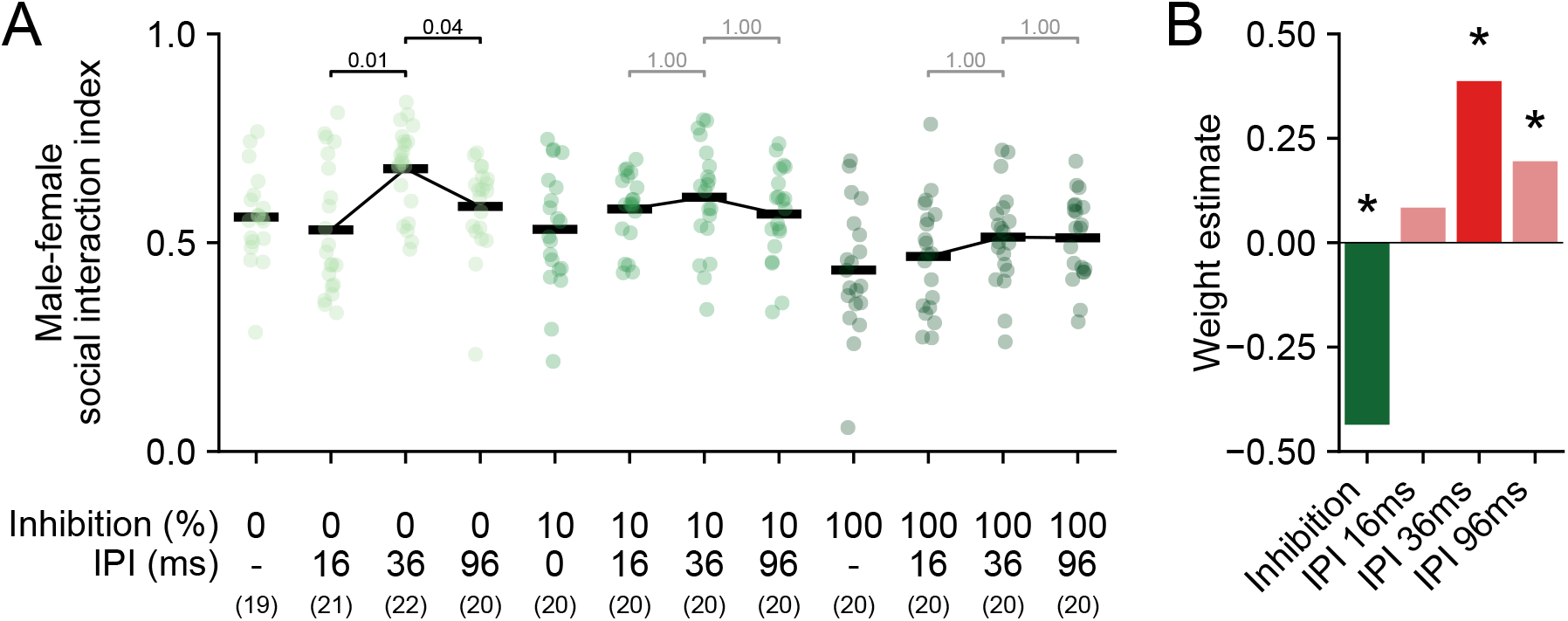
The mode of integration for song and female taste cues to control female-directed social interactions is independent of inter-pulse interval (IPI). **A** Social interaction index for different levels of taste inhibition and IPIs. Without inhibition (0%, lightest green), the conspecific IPI of 36 ms had a stronger effect than shorter or longer IPIs. Increasing inhibition flattened the IPI tuning. **B** Weights from the selected GLM (linear vs. null: *p* = *2 .0 e*^*−10*^, nonlinear vs. linear: *p* = *0 .38*) predicting the female-directed social interactions from A (see tables S5 and S6 for details). Inhibition of female taste cues is associated with a negative weight (*w*_*inh*_ = *− 0 .44, p* = *2 .1 e*^*−9*^, green) indicating that interference reduced social interactions. Weights for the different IPIs were highest for the conspecific IPI (36 ms, *w* = *0 .39, p* = *2 .4* ^*−5*^, red), weaker for the longer IPI (96 ms, *w* = *0 .20, p* = *0 .03*, light red), and non-significant for the shortest IPI (16 ms, *w* = *0 .08, p* = *0 .36*, light red). Asterisk above bar indicate significance of weight in the model. Dots in A correspond to individual male-male pairs (N provided below the panel). Green darkness in A encode the inhibition strength, as indicated below each group. For pair-wise comparisons, we tested involved groups for normality with a Shapiro test, and conducted the comparison with an independent t-tests for unequal variances if both groups were normally distributed, or with a two-sided Mann-Whitney U test otherwise (first three pair comparisons from left to right). All p-values are reported after Bonferroni correction for multiple comparisons.

Linear integration was sufficient to explain the data (Fig. 4B). Inhibition of ppk23 neurons was associated to a strong negative weight (*w*_*inh*_ = *− 0 .44, p* = *2 .1 e*^*− 9*^). The IPI weights were highest for the conspecific IPI (36 ms, *w* = *0 .39, p* = *2 .4 e*^*−5*^), lower for the 96 ms (*w* = *0 .20, p* = *0 .03*), and non-significant for the 16 ms (*w* = *0 .08, p* = *0 .36*). These results show that the observed linear song-taste integration is independent and selectively tuned to the conspecific IPI.

### Song and female taste are integrated nonlinearly to control courtship

Finally, we investigated how song influences a male’s decision to engage in courtship or aggression when interacting with females. As shown above, song increases aggression towards males, independent of taste cues (Fig. 2A). However, males never displayed aggression toward females, regardless of manipulations (Fig. 5A). This suggests that song is not sufficient to drive female-directed aggression and that female taste cues do not gate aggression.

**Figure 5:**
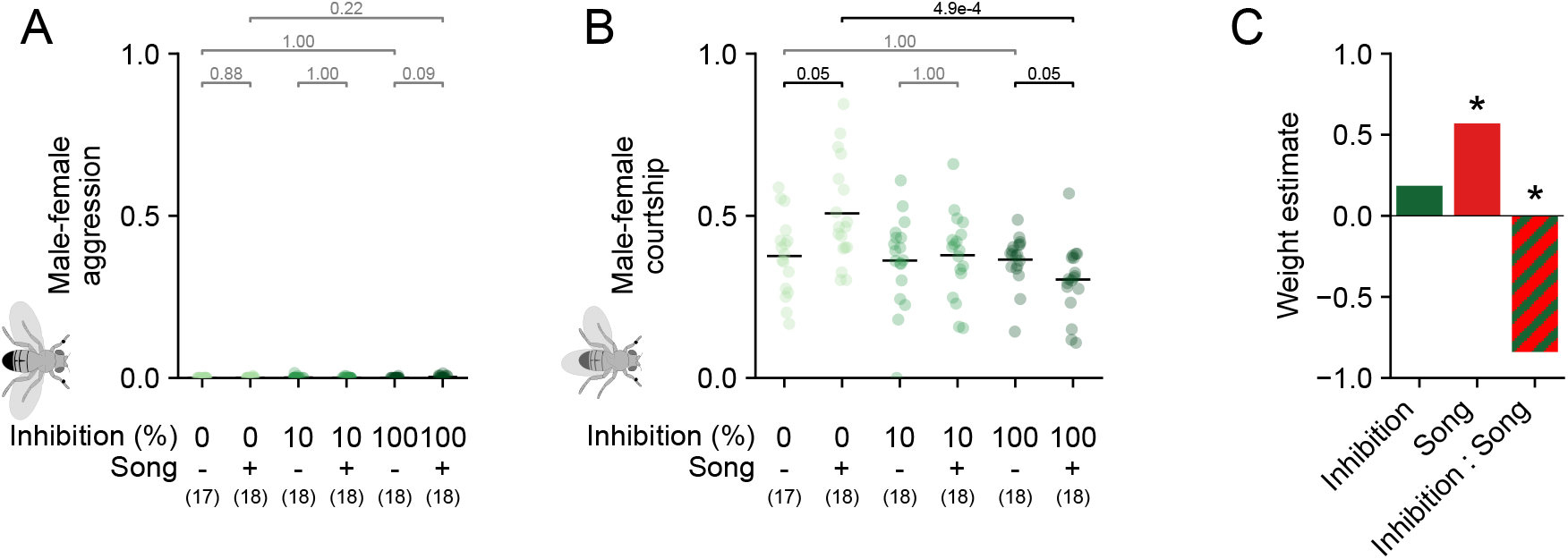
Female taste cues flip the valence of courtship song. **A** Probability of aggression during interactions. Model fit omitted due to lack of significant effects. **B** Probability of courtship during female-directed social interactions. **C** Weights from the selected GLM (linear vs. null: *p* = *6 .0 e*^*−3*^, nonlinear vs. linear: *p* = *0 .05*) predicting the courtship probabilities shown in B (see tables S3 and S4 for details). Inhibition of female taste cues alone was not significant (*w*_*inhibition*_ = *0 .18, p* = *0 .39*, green). Song is associated with a positive weight (*w*_*song*_ = *0 .57, p* = *1 .0 e*^*−3*^, red) and promotes display of courtship. Interaction of inhibition and song resulted in a significant, strong negative weight (*w*_*inhibition*:*song*_ = *− 0 .84, p* = *5 .0 e*^*−3*^, red-green stripes), which flips the valence of song to suppressing courtship when female taste cues are inhibited. Asterisk above bar indicate significance of weight in the model. Dots in A and B correspond to individual male-male pairs (N provided below the panel). Green darkness in A and B encode the inhibition strength, as indicated below each group. For pair-wise comparisons, we tested involved groups for normality with a Shapiro test, and conducted the comparison with an independent t-tests for unequal variances if both groups were normally distributed (comparisons involving inhibition levels 0% and 10% in B), or with a two-sided Mann-Whitney U test otherwise. All p-values are reported after Bonferroni correction for multiple comparisons.

**Figure 6:**
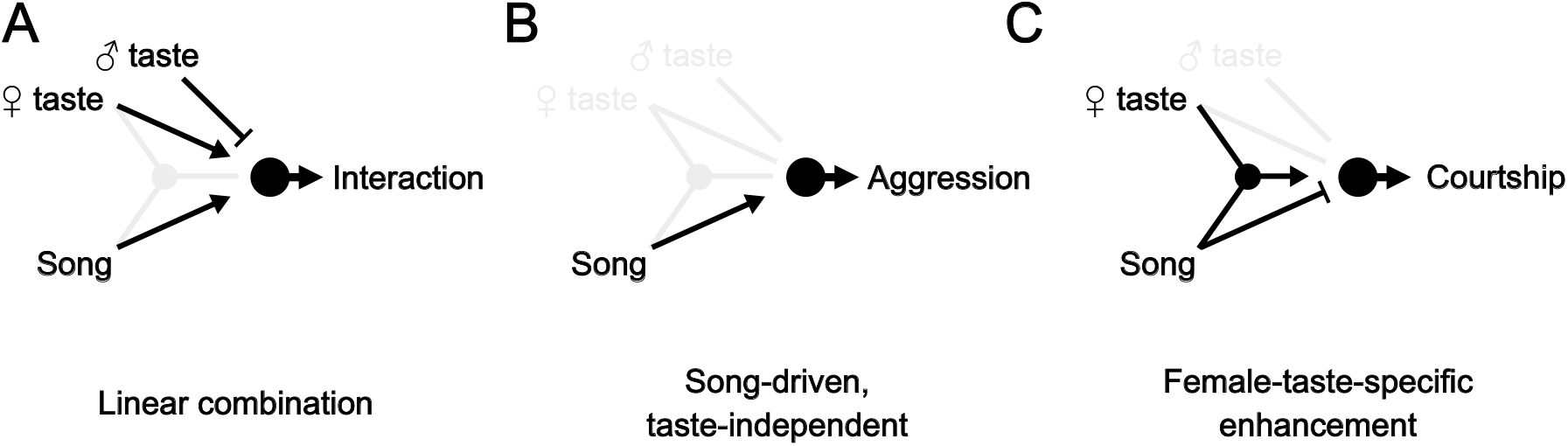
Males integrate taste and song with three context-specific strategies during social behavior. **A** Males linearly integrate male and female taste with courtship song in the modulation of general social interaction: male taste cues suppress (blunt-ended arrow), while female taste cues and song promote (pointed arrow) interactions. Effects with not contributions, in this case the nonlinear combination of song and taste, are colored in gray. **B** Song promotes aggression towards males, with no effect from male or female taste cues. **C** Song and female taste cues are nonlinearly integrated to control courtship towards females. Song alone weakly suppresses courtship. Song in the presence of female taste cues promotes courtship towards females.

The effect of song on courtship toward females strongly depended on the males’ ability to perceive female taste cues (Fig. 5B). When males were able to fully perceive female taste cues (0% inhibition), song promoted courtship. Weak optogenetic inhibition of the taste receptors in the foreleg tarsi, abolished the courtship-promoting effect of song. Surprisingly, strong inhibition inverted the valence of song from promoting to suppressing courtship. This switch in the valence of song with female taste cues indicates a nonlinear song-taste integration (Fig. 5B). Accordingly, a nonlinear integration model best explained female-directed courtship (Fig. 5C). Song had a strong positive effect on courtship (*w*_*s*_ = *0 .57, p* = *1 .0 e*^*− 3*^, red), while inhibition of taste was not a significant predictor (*w*_*inh*_ = *0 .19, p* = *0 .39*, green; Table S4). However, a negative interaction term (*w*_*inh*_ = *− 0 .84, p* = *5 .2 e*^*− 3*^, red and green stripes) indicates that the effect of song depends on the presence of female taste cues. Since the interaction term is opposite in sign and larger in magnitude than the effect from song, it not only reduces but reverses the effect of song when the detection of female taste cues is inhibited.

## Discussion

We show that male *Drosophila melanogaster* integrate courtship song and taste cues using three context-specific integration strategies. First, when deciding whether to interact with another fly, males integrate cues linearly: female taste and song increase interaction, while male taste independently suppresses interaction (Fig. 1D-E and 3B-C). Second, when deciding whether to be aggressive towards another male, males rely only on song, and ignore taste cues (Fig. 2A). Third, when deciding whether to court a female, song and taste are nonlinearly integrated: Female taste cues switch the behavioral valence of song, increasing courtship (Fig. 5B).

To interpret these diverse strategies, we consider that cue combinations signal distinct types of information and behavioral contexts: Taste cues are short-range and reliably signal the sex of the frontal fly (Clowney et al., 2015; Kallman et al., 2015; Seeholzer et al., 2018). By contrast, courtship song is more long range than taste and originates from another male singing to a female. Song thus signals a courtship context and the presence of at least one male competitor and female (Schilcher, 1976; Yoon et al., 2013; Li et al., 2018; Hindmarsh Sten et al., 2025).

When deciding whether to interact socially, males integrated song and taste cues linearly (Fig. 1E, 3C), suggesting that these modalities provide largely independent bits of information. Hearing song signals that a female is being courted by another male, and should therefore arouse the male to prepare him for courtship and aggression. Independent of that, male taste suppresses, while female taste boosts interactions, to prioritize female-directed courtship over other types of interactions.

When deciding whether to be aggressive, males only relied on song and ignored male taste cues (Fig. 2B). This likely occurs because song reliably signals the presence of a male competitor, which should drive aggression regardless of whether the frontal fly is recognized as a male. A female target never elicited aggression, even when the male was unable to detect taste cues, because 11-cis-vaccenyl acetate (cVA), a male specific olfactory cue is, required to drive aggression (Hindmarsh Sten et al., 2025).

When deciding whether to court, males nonlinearly integrated song and taste: In the presence of female taste cues, song increased courtship, but in their absence, song weakly suppressed courtship (Fig. 5C). This reflects how female taste cues signal different outcomes of a courtship competition: Hearing song while detecting female taste cues indicates that the frontal fly is the female, with a competitor nearby. In this context, males should increase courtship to elicit receptivity (vaginal plate opening) and attempt mating before loosing access to the female (Wang et al., 2021; Mezzera et al., 2020). By contrast, hearing song without female taste cues implies that the sex of the frontal fly is unclear and that the courter does not have access to the female. In this context, courtship is futile. Instead, the male should be more aggressive to gain access to the female.

Thus, these diverse, context-specific integration strategies are consistent with principles of efficient multimodal integration in which reliable cues are upweighted and irrelevant cues are suppressed (Alais and Burr, 2004; Körding et al., 2007; Cao et al., 2019). They also show how males can make better decisions by considering multiple cues, e.g. when female taste cues modulate the effect of song to prevent futile courtship or aggression towards the female.

Song has been previously shown to trigger courtship in groups of males (Yoon et al., 2013; Li et al., 2018; Zhou et al., 2015), but not in our experiments with pairs of males (Fig 2B). This suggests that the group context, with an increased density of social cues, is crucial for song-driven chaining. This may be explained by amplified motion cues, which alone can drive male courtship (Kohatsu and Yamamoto, 2015; Ribeiro et al., 2018; Hindmarsh Sten et al., 2021) and which are further increased in a group by song promoting locomotion in males (Crossley et al., 1995; Deutsch et al., 2019).

How are these diverse behavioral strategies for song-taste integration implemented in the brain? While our results do not identify the mechanisms underlying song-taste integration, we can hypothesize about possible neural substrate underlying the three strategies. Song-taste integration could occur early, near the sensory periphery (Iurilli et al., 2012; Ibrahim et al., 2016; Teichert and Bolz, 2017; Teichert and Bolz, 2018), or late, in central integration or decision centers (Coen et al., 2023; Rohe and Noppeney, 2016; Cao et al., 2019).

The connectome reveals abundant connections between early sensory neurons from multiple modalities (Walker et al., 2025; Schlegel et al., 2024; Dorkenwald et al., 2024; Stürner et al., n.d.), providing substrates for early multisensory integration. However, early integration can support diverse modes of integration only if song is processed in parallel pathways. This possibility is supported by redundant representations of song in the fly brain (Baker et al., 2022). For example, vPN1, pC2l, and CL062 (called AVLP_pr12 in Baker et al. (2022)) neurons detect pulse song and drive distinct behaviors (Zhou et al., 2015; Deutsch et al., 2019; Nair et al., 2025; Tao et al., 2024). Their responses may be differentially modulated by pheromones to implement the distinct multimodal integration strategies observed in our study. Similarly, male and female taste cues are processed in parallel pathways, which may be differentially modulated by song. For instance, vAB3 and PPN1 both encode female-specific taste cues but project to different downstream targets (Clowney et al., 2015; Kallman et al., 2015).

Multimodal integration could also occur centrally. Like a switchboard, parallel decision centers could flexibly link sensory cues with behavior, depending on the combinatorial activation from multimodal sensory cues (Schretter et al., 2025; Hindmarsh Sten et al., 2025). In *Drosophila*, different subtypes of the pC1 neurons drive distinct social behaviors depending on the sensory context (Koganezawa et al., 2016; Hindmarsh Sten et al., 2025; Hoopfer et al., 2015; Hoopfer, 2016; Zhou et al., 2014; Zhou et al., 2015): The pC1x neurons drive aggression in males. These neurons are activated by song and olfactory cues (Hindmarsh Sten et al., 2025), but have not been shown to respond to gustatory cues. This aligns with our behavioral finding that aggression is driven by song, but not by taste cues, and requires cVA from a nearby male (Fig. 2A, 5A) (Kurtovic et al., 2007; Wang and Anderson, 2010; Hindmarsh Sten et al., 2025). This suggests that pC1x neurons could be a locus of taste-independent, song-driven aggression directed at males. The P1a neurons encode a persistent social arousal state and drive courtship and aggression behaviors in a context-dependent manner (Hoopfer et al., 2015; Jung et al., 2020). They are activated by female taste cues and suppressed by male taste and olfactory cues (Clowney et al., 2015; Kallman et al., 2015). This is consistent with our findings that male taste cues suppress, and female taste cues promote, social interactions (Fig. 1, 3). P1a neurons respond to song only during visually-induced courtship (Hindmarsh Sten et al., 2025). This suggests that song activates P1a only when a male has been primed to court by female taste cues, consistent with our observation that song promotes courtship only in the presence of female taste cues. However, how song and taste cues are combined in P1a remains unknown. The pCd neurons are downstream of P1a, are activated by male olfactory cues, and are required for persistent male courtship and aggression (Zhang et al., 2018; Jung et al., 2020). Activation of P1a enhances pCd responsiveness to male chemical cues. Similar modulation may occur with female chemical cues, potentially enabling the nonlinear integration of song and female taste cues. Multimodal integration could also emerge from changes in neuronal dynamics, with male or female cues enhancing the persistence of pCd activity, and thus courtship or aggression.

Finally, different multimodal integration strategies may be implemented in pre-motor circuits by specific cues gating context-specific descending pathways (Cande et al., 2018; Braun et al., 2024). Whether P1a activation drives courtship or aggression depends on chemical cues, with male cues promoting aggression and female cues promoting courtship (Hoopfer et al., 2015; Jung et al., 2020). Therefore, nonlinear song-taste integration may result from female taste cues facilitating or gating song-induced courtship.

Overall, this suggests how the modular architecture of the brain could support flexible, context-specific integration strategies. Our finding that males use diverse strategies to integrate song, taste and other cues sets the stage for identifying the circuit mechanisms by which multimodal integration is implemented in the brain.

## Methods

### Fly strains and husbandry

Flies were maintained on a 12 h light:12 h dark cycle at *25 °C* and 60% humidity. Males and females were anesthetized with carbon dioxide and sorted within 2 hours after eclosion. Males were individually housed in food vials to ensure social isolation, while females were grouped in cohorts of 5–10 flies per vial. All experiments were conducted on flies within their 4-7 days after eclosion, at approximately *25 °C*, and within the first 3 hours of their circadian cycle. Wild-type assays were performed with males and females of the *Drosophila melanogaster* CantonS-Tully strain, referred to as *Canton S* in the text. All females used for the experiments in this study were *Canton S* virgin females, 4-7 days old, deafened by cutting their aristae. Optogenetic assays were performed with heterozygous males with genotypes *w/+* ; *ppk23-GAL4/+* ; *20XUAS-GtACR1 (attP2)/+* from the progeny of ; *ppk23-GAL4* ; (Clowney et al., 2015; Kallman et al., 2015) and *w* ; ; *20XUAS-GtACR1 (attP2)* (Mauss et al., 2017). These males are referred to as *ppk23+-GtACR1*. Their parental lines are referred to as *ppk23-Gal4* and *UAS-GtACR1*.

### Behavioral experiments

Experiments were conducted in sound-isolated boxes, each equipped with a Raspberry Pi 4 (Model B 4 GB RAM) for stimulus control and video recording. Stimuli were presented via a Hi-Vi Research® loudspeaker Model F6 and a LED array (see Optogenetics section in Methods). Videos were recorded using a Raspberry Pi High-Quality Camera (12.3 MP, 7.9 mm diagonal) at 1232×1640 pixels and 30 Hz. All experiments were performed with infrared illumination from the back of the behavioral arenas (12 V LED Strip IR 850 - IR 1-60-850, SOLAROX®).

Each behavioral arena was 3 mm high and 18 mm in diameter. The experimental setup consisted of holders with 10 behavioral arenas per experiment. Each arena was divided into two sections by a movable barrier. The opaque walls and transparent ceiling of the arenas were polished to diminish the amount of time spent on the walls or ceiling. The floor consisted of a fine mesh that allowed sound and infrared light to pass through the arena.

Flies were gently aspirated into each side of the movable barrier, separating the male fly from their corresponding target. Once all flies were loaded into the arenas, the holder was carefully placed inside the isolation box, and the barriers were opened at the start of the recording.

### Optogenetics

For optogenetic experiments, male flies were housed on fly food containing retinal (1 ml Sigma-Aldrich all-trans retinal solution per 100ml of food, dissolving 100 mM in 95% ethanol). Food vials were wrapped in aluminum foil to prevent degradation and unintended activation of optogenetic channels.

GtACR1 channels were activated using an array of eight green LEDs (525 nm wavelength, WINGER® WEPRGB9-S1 Power LED Star RGB 9 W) homogeneously distributed above the behavioral arenas. The three green LED light intensities used during the experiments are represented as percentages in the figures and correspond to the following measurements: *0 W* /*m*^*2*^ (0%), *0 .313 W* /*m*^*2*^ (10%), and *2 .500 W* /*m*^*2*^ (100%). All optogenetic experiments were conducted under dim white light to preserve visual cues without significantly activating the GtACR1 channels.

### Courtship song playback

The song stimuli consisted of an artificially generated pulse song. The stimulus block consisted of 20 s of song interleaved with 120 s of silence, and the stimulus block was repeated for the duration of the experiment (30 min). The song stimulus was generated as in Deutsch et al. (2019), with a sampling rate of 10 kHz. Gabor wavelets (16 ms duration) were arranged to produce the desired interpulse interval (e.g., a 20 ms pause for a 36 ms IPI). Gabor wavelets were generated by modulating the amplitude of a short sinusoidal wave using a Gaussian: exp(*− t*^*2*^ /(*2 σ*^*2*^)) sin(*2 πf ∗ t* + *ϕ*), where *f* is the pulse carrier frequency, *ϕ* is the phase carrier, and *σ* is proportional to the pulse duration. For all stimuli in this study, we used a pulse carrier frequency of 250 Hz and pulse carrier phase of *π*/*4*.

### Manipulations

All manipulations were performed between 18–24 hrs prior to the experiment. We used cold temperature to anesthetize the flies prior to the manipulations and kept the flies on a cold block of aluminum during the manipulation, which lasted approximately 20 s per individual. Flies recovered in inverted food vials to prevent adherence to the food surface. They were returned to the incubator after regaining normal mobility.

To remove the foreleg tarsi, males were positioned on their backs, with their legs up, and the foreleg tarsi were carefully clipped with forceps at the position of the sex comb.

To control for any effects induced through the manipulation, *Canton S* males that did not undergo tarsi removal were still ice anesthetized and placed on a cold block of aluminum for approximately 20 s, and left to recover.

To render the females deaf, they were positioned on their backs, and we carefully cut their aristae with *3 mm Vannas Spring Scissors* (Fine Science Tools No. 15000-00). To consider the manipulation successful, the cut had to be sufficiently close to the antennae to leave only a stub with no bristles.

### Fly tracking

We used SLEAP (Pereira et al., 2022) for tracking the position of the flies in the *Canton S* and *ppk23+-GtACR1* assays. We used the resulting tracks to calculate the distance between the flies and their velocities.

### Social interaction index

We calculated the social interaction index as the fraction of time a pair of flies was closer than 4 mm. To exclude periods when males were idle next to each other without actively interacting, we also required that at least one of the flies was moving faster than 1 mm/s. To control for outliers from tracking errors or fitness-related factors, we excluded pairs with more than 50% tracking errors (one case per assay) or those exhibiting extremely low social interaction indices (less than 0.05; three cases in *Canton S*, four in *ppk23+-GtACR1*).

### Manual annotations

Video recordings were of insufficient quality to track the wings. We therefore manually annotate unilateral and bilateral wing extensions. Using DAS (Steinfath et al., 2021), we annotated frames 8 s apart as containing unilateral, bilateral, or no wing extension. Instances of grooming or attempts of flying were excluded. To reduce bias, annotators were blind to the conditions of the experiments.

### Generalized linear models

Generalized linear models (GLMs) were fitted in R with the glmmTMB package. All independent variables were coded as categorical variables. To fit the GLMs, we selected the family function beta_family, which uses a logit link function. Model comparisons were performed using glmmTMB. The primary approach used a *χ*^*2*^ test (whether *p < 0 .05*) to assess whether more complex models significantly improved predictions (“linear vs null” and “nonlinear vs linear”). The second approach was based on the Akaike Information Criterion (AIC) (Akaike, 1974), which measures the relative amount of information lost by a model for a given set of data, penalizing for an increased number of parameters in the model, thus providing a fair comparison across models with different numbers of parameters. Confidence intervals were estimated via bootstrapping, using custom R code provided by Roger Mundry. The model *p-values* were calculated through comparison of models with inclusion and omission of a given term (for more detail, see documentation of glmmTMB).

For the models involving *Canton S* flies, only a subset of the experiments was annotated for courtship and aggression, since the required manual annotations were time consuming. Thus, GLMs for social interaction index used the complete dataset (*n ≈ 30*), while courtship and aggression analysis were done with the annotated subset (*n ≈ 20*). For the models involving optogenetic inhibition, we used the min-max scaled log(*1* + *LE D*) to account for the nonlinearity between light and neuron inhibition.

### Software and packages

**Table 1:** Software and packages.

| Resource | Identifier |
| --- | --- |
| SLEAP | https://sleap.ai (Pereira et al., 2022) |
| scikit-learn | https://scikit-learn.org (Pedregosa et al., 2011) |
| python (>=3.9) | https://python.org |
| R (4.2.3) | https://www.R-project.org/ (R Core Team, 2021) |
| DAS | https://github.com/janclemenslab/das (Steinfath et al., 2021) |
| statmodels | https://www.statsmodels.org/ (Seabold and Perktold, 2010) |
| seaborn | https://seaborn.pydata.org/ (Waskom, 2021) |
| scipy | https://scipy.org/ (Virtanen et al., 2020) |
| matplotlib | https://matplotlib.org/ (Hunter, 2007) |
| glmmTMB | https://cran.r-project.org/package=glmmTMB/ (Brooks et al., 2017) |

## Author contributions

- Conceptualization, Writing - APM, JC
- Investigation, Data curation, Formal analysis - APM

## Acknowledgments

We thank Roger Mundry for advice on GLM modeling. This study was funded by an ERC Starting Grant NeuSoSen (851210) and a DFG Emmy Noether Grant (329518246) to JC.

## Supplemental Information

**Table S1:**
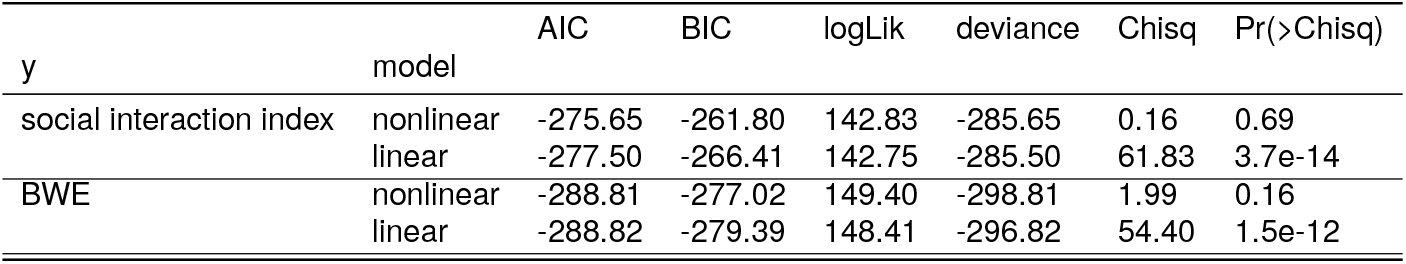
Canton S male-male model selection. Relevant to Fig. 1, 2 and 3.

**Table S2:** Canton S male-male best model results. Related to Fig. 1, 2 and 3.

| y | model | name | Estimate | Std. Error | z value | Pr(>\|z\|) |
| --- | --- | --- | --- | --- | --- | --- |
| social interaction index | linear | bias | -1.17 | 0.07 | -18.01 | 1.6e-72 |
|  |  | taste | -0.47 | 0.08 | -6.28 | 3.5e-10 |
|  |  | song | 0.47 | 0.07 | 6.26 | 3.8e-10 |
| BWE | linear | bias | -3.27 | 0.22 | -15.18 | 4.5e-52 |
|  |  | taste | -0.02 | 0.19 | -0.10 | 0.92 |
|  |  | song | 1.78 | 0.22 | 8.16 | 3.5e-16 |

**Table S3:** ppk23-GtACR1 conspecific song playback model selection. Related to Fig. 3 and 5.

| y | model | AIC | BIC | logLik | deviance | Chisq | Pr(>Chisq) |
| --- | --- | --- | --- | --- | --- | --- | --- |
| social interaction index | nonlinear | -148.25 | -134.27 | 79.12 | -158.25 | 0.11 | 0.74 |
|  | linear | -150.14 | -138.95 | 79.07 | -158.14 | 37.58 | 6.9e-09 |
| UWE | nonlinear | -85.76 | -72.39 | 47.88 | -95.76 | 7.55 | 6.0e-03 |
|  | linear | -80.20 | -69.51 | 44.10 | -88.20 | 5.82 | 0.05 |

**Table S4:** ppk23-GtACR1 conspecific song playback results. Related to Fig. 3 and 5.

| y | model | name | Estimate | Std. Error | z value | Pr(>\|z\|) |
| --- | --- | --- | --- | --- | --- | --- |
| social interaction index | linear | bias | 0.23 | 0.08 | 2.93 | 3.4e-03 |
|  |  | inhibition | -0.56 | 0.11 | -5.18 | 2.2e-07 |
|  |  | song | 0.38 | 0.10 | 4.02 | 5.7e-05 |
| UWE | nonlinear | bias | -0.72 | 0.13 | -5.71 | 1.1e-08 |
|  |  | inhibition | 0.18 | 0.21 | 0.87 | 0.39 |
|  |  | song | 0.57 | 0.17 | 3.29 | 1.0e-03 |
|  |  | inhibition : song | -0.84 | 0.30 | -2.79 | 5.0e-03 |

**Table S5:** ppk23-GtACR1 model selection, including all IPIs. Related to Fig. 4.

| y | model | AIC | BIC | logLik | deviance | Chisq | Pr(>Chisq) |
| --- | --- | --- | --- | --- | --- | --- | --- |
| social interaction index | nonlinear | -315.04 | -283.64 | 166.52 | -333.04 | 3.11 | 0.38 |
|  | linear | -317.93 | -297.00 | 164.97 | -329.93 | 51.22 | 2.0e-10 |

**Table S6:** ppk23-GtACR1 GLM results, including all IPIs. Related to Fig. 4.

| y | model | name | Estimate | Std. Error | z value | Pr(>\|z\|) |
| --- | --- | --- | --- | --- | --- | --- |
| social interaction index | linear | bias | 0.19 | 0.07 | 2.63 | 8.0e-03 |
|  |  | inhibition | -0.44 | 0.07 | -5.99 | 2.1e-09 |
|  |  | IPI 16ms | 0.08 | 0.09 | 0.92 | 0.36 |
|  |  | IPI 36ms | 0.39 | 0.09 | 4.23 | 2.4e-05 |
|  |  | IPI 96ms | 0.20 | 0.09 | 2.12 | 0.03 |

